# Data-independent acquisition (DIA) approach for comprehensive ubiquitinome profiling in targeted protein degradation

**DOI:** 10.1101/2025.05.27.656291

**Authors:** Gajanan Sathe, Sascha Röth, Géraldine Gelders, Abigail Brewer, Thomas J. Macartney, Nicola T Wood, Louis De Muynck, Mark A Nakasone, Toan K Phung, Arjan Buist, Diederik Moechars, Gopal P. Sapkota

## Abstract

Targeted protein degradation (TPD) has emerged as a highly promising therapeutic strategy for a wide range of diseases, including cancer and neurodegenerative disorders. The ubiquitin-proteasome system, which is responsible for protein degradation, plays a critical role in this process. Gaining comprehensive insights into the ubiquitylation landscape is essential for the development of selective and efficient targeted protein degradation approaches. Recently, data-independent acquisition (DIA) has gained significant popularity as a robust and unbiased approach for quantitative proteomics. Here, we report a cutting-edge workflow that utilizes diGly antibody-based enrichment followed by an optimized Orbitrap-based DIA method for the identification of ubiquitylated peptides. We identify over 40,000 diGly precursors corresponding to more than 7,000 proteins in a single measurement from cells exposed to a proteasome inhibitor, highlighting an exceptional throughput. By applying our optimized workflow, we successfully identify ubiquitylation sites on substrate proteins with various TPD approaches. Therefore, our workflow holds tremendous potential for rapidly establishing mode of action for various TPD modalities, including PROTACs and molecular glues.

## Introduction

Protein stability exhibits a dynamic range within cells, with some proteins having longer half-lives than others. One crucial factor influencing this stability is post-translational modification (PTM), specifically ubiquitylation, which plays a fundamental role in virtually all cellular processes, including protein turnover ^1^. The ubiquitylation process involves a cascade of events, encompassing a ubiquitin-activating enzyme (E1), a ubiquitin-conjugating enzyme (E2), and a ubiquitin-ligating enzyme (E3). This cascade culminates in the covalent attachment of a 76-amino acid ubiquitin molecule to primarily a lysine residue’s ε-amine group on the substrate protein ^2,3^. The reversal of this modification is orchestrated by a family of enzymes known as deubiquitinating enzymes (DUB) ^4^. Further ubiquitin molecules can be covalently added onto the N-terminus or the ε-amine groups of one or more of the seven lysine residues of the ubiquitin molecule already on the substrate, resulting in polyubiquitin chains of various architectures. Different chain configurations can encode distinct biological functions. For example, Lys48-linked chains, which are the most common type, target proteins for proteasomal degradation ^5^. Lys63-linked chains, the second most prevalent, encode non-degradative functions ^6^. Emerging research has revealed various biological roles of other ‘atypical’ polyubiquitin chains, such as Met1, Lys6, Lys11, Lys27, Lys29, or Lys33 linkages ^7^. Similarly, diverse roles for branched polyubiquitin chains have also been reported^8,9^. In normal cell physiology, the process of reversible ubiquitylation is tightly regulated. Therefore, dysregulation of ubiquitylation events have been associated with various diseases, such as neurodegenerative disorders ^10^, autoimmune conditions ^11^, and inflammatory diseases ^12^.

Initial advances in ubiquitin research relied on methods involving overexpression of tagged ubiquitin, enrichment with affinity tag binders or ubiquitin-specific antibodies, and, more recently, ubiquitin-binding traps to investigate the role and extent of ubiquitylation in cells. A major limitation of these methods was the difficulty in mapping sites of ubiquitin modification. Although tandem ubiquitin binding entities (TUBEs) offer a good enrichment strategy, direct mapping of ubiquitylated lysine residues on a global scale is still not feasible ^13^. More recently, two different antibody approaches have transformed our ability to enrich and identify ubiquitylated peptides. During protein trypsinization, ubiquitylated proteins generate a signature diGly remnant that can be recognised and enriched by a specific antibody that recognises the diGly remnant peptides ^14^. Several studies have used this approach to enrich diGly remnant peptides, which can then be identified by mass-spectrometry ^15,16^. Other antibodies, such as the UbiSite antibody which recognises the C-terminus of ubiquitin, have also been developed to enrich and identify precise sites of ubiquitylation ^17,18^. To make it more high-throughput and quantitative, DiGly enrichment approach has been combined with tandem mass tag (TMT) labelling ^19^, and on-antibody TMT labelling ^20^, for data-dependent acquisition (DDA) approaches to global ubiquitinome profiling. In recent years, data-independent analysis (DIA) has gained significant traction as a robust and unbiased approach for quantitative proteomics. This method allows for comprehensive proteome profiling by systematically fragmenting and quantifying all detectable peptides within a given sample ^21,22^. Several high throughput studies have utilised DIA workflows for the identification of the ubiquitinome and reported higher number of ubiquitylated peptides compared to the DDA ^23,24^. There has also been a significant advancement of the analytical platforms to evaluate DIA data using MaxDIA ^25^, Spectronaut ^26^, and DIA-NN ^27^. These analytical workflows were also used for global DiGly ubiquitinome mapping, which have reported the highest number of global DiGly peptides to date ^23,28^.

Targeted protein degradation (TPD) approaches harness cellular ubiquitin-proteasome or autophagic pathways to induce the destruction of specific proteins of interest (POIs). To glean mechanistic insights into and the specificity of POI degradation, it is pertinent to map the precise ubiquitylation sites on the POI as well as global changes in ubiquitylation caused by the degrader. By leveraging the advances in mapping ubiquitylation sites, we have developed cutting-edge workflows that combine diGly antibody-based enrichment with an optimized Orbitrap-based DIA method for the identification of ubiquitylated peptides in TPD. Our workflows provide a comprehensive and detailed overview of the ubiquitin landscape within the cellular proteome. We have successfully adopted and adapted these workflows to uncover POI ubiquitylation sites for several TPD approaches, including the affinity-directed protein missile (AdPROM) system ^29^, dTAG-degron knockin on target POI ^30^, and for a proteolysis targeting chimera (PROTAC) targeting the degradation of an endogenous protein ^31^.

## Results

### Optimization of the cell lysis methods for the mass spectrometry-based analysis of global ubiquitylation

We compared two well-established cell lysis methods that are frequently employed in proteomics, namely urea-based and SDS-based, for the identification of diGly peptides. After treating HEK293 cells with the proteasome inhibitor MG-132 (20 µM) for 6 h, cells were lysed with each lysis method. For the urea-lysis method, cells were lysed in a buffer containing 8 M urea and sonicated, and 2 mg protein from extracts subjected to in-solution digestion ^32^. The digested peptides were cleaned using C_18_ Sep-Pak columns (Fig. 1A). For the SDS-based lysis, we followed the S-Trap™ sample preparation protocol ^33,34^, where cells were lysed in a buffer containing 2% SDS, extracts heated to 95°C for 5 min and then sonicated. 2 mg protein from extracts were subjected to in-solution digestion and peptides were then cleaned (Fig. 1A). The cleaned peptides were subjected to diGly-remnant enrichment using the anti-diGly antibody conjugated to agarose beads. The eluted diGly-peptides were analysed using an Orbitrap Exploris 480 mass spectrometer using data-dependent acquisition (DDA). From both lysis methods, DDA analysis identified a total of 20,941 diGly-modified peptides (Fig. 1B&C) corresponding to 4,638 proteins (Fig. 1D&E). Among these, the urea-based lysis method identified a total of 15,868 DiGly peptides corresponding to 4,092 proteins from 3 replicates. Each replicate identified over 9000 diGly peptides, with a coefficient of variation (CV) <20% (Fig. 1B&D; Supp. Data 1). The SDS-lysis method identified a total of 11,820 diGly peptides corresponding to 3,436 proteins from 3 replicates. Each replicate contained over 6,000 DiGly peptides, with a CV<20% (Fig. 1B&D; Supp. Data 1). By comparison, the urea-lysis method identified 34% more diGly peptides (Fig. 1B) and 16% more ubiquitylated proteins (Fig. 1D) compared to the SDS-lysis workflow. 6747 diGly peptides (Fig. 1C) corresponding to 2890 proteins (Fig. 1E) were identified by both lysis methods.

**Figure 1.**
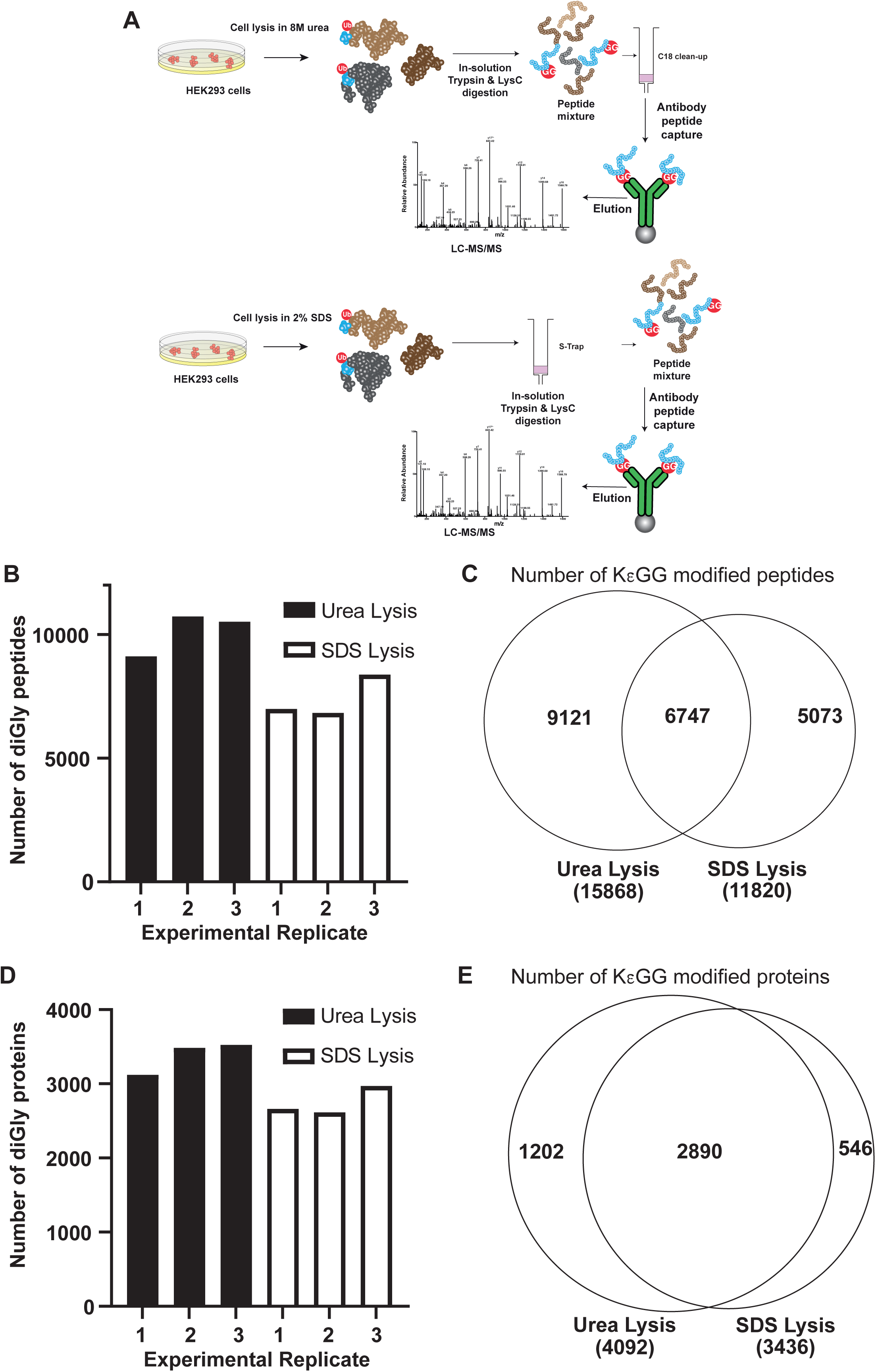
Comparison of cell lysis methods for in-depth ubiquitinome analysis. **(A)** Schematic illustrating the two routinely employed cell lysis methods used for diGly ubiquitinome profiling by mass spectrometry: 8 M urea lysis and 2% SDS lysis. The cartoons illustrate an overview of key steps used for each lysis method. **(B)** Bar chart showing the number of diGly peptides identified using each of the lysis methods from (A) from each of the experimental replicates. **(C)** Venn diagram showing the number of common and unique diGly peptides identified using the 8M urea or 2% SDS lysis buffer systems. **(D)** As in (B), except the chart shows the numbers of diGly modified proteins identified. **(E)** As in (C), except the Venn diagrams show the numbers of diGly modified proteins identified.

### Comparison between data-dependant acquisition (DDA) and data-independent acquisition (DIA) methods for global ubiquitylation analysis

To optimize the mass spectrometry workflow for a comprehensive analysis of global ubiquitylation, we conducted a comparative analysis of DDA and DIA methods. For this, we took the urea-lysis method that yielded higher numbers of diGly peptides (Fig. 1). By subjecting 2 mg protein from HEK293 extracts for digestion followed by a diGly enrichment, the diGly-enriched peptides were analysed on Orbitrap Exploris 480 mass spectrometer equipped with both DIA and DDA modes. We optimized the MS method for our analysis with a 120-minute LC gradient (Supp. Data 2). We then benchmarked the DIA workflow against the state-of-the-art DDA approach described above (Fig. 1). For DIA data processing, we employed DIA-NN using library-free mode, whereby we performed searches against a sequence database without relying on an experimentally generated spectral library. Conversely, for DDA data processing, we utilized the SEQUEST search engine ^35^. This comprehensive evaluation allowed us to assess the performance of both DIA and DDA workflows in the context of global ubiquitylation analysis. In each of the 3 replicates, the DDA analysis identified 9426, 10834, 11166 unique diGly peptides corresponding to 3122, 3487, 3525 proteins, respectively (Fig. 2A&B, Supp. Data 1). On the other hand, DIA analysis identified 25775, 33400, 28195 diGly peptide spectral matches corresponding to 5291, 6140, 5522 proteins, respectively (Fig. 2A&C, Supplementary Data 3&4). By direct comparison, DIA identified a total of 38883 diGly peptides corresponding to 6646 proteins, while DDA identified 15868 diGly peptides corresponding to 4092 proteins (Fig. 2A&C). In a direct comparison of DiGly-modified sites, DDA identified 13135 sites, while DIA identified 32141 sites across three replicates. Of these, 9573 sites were detected by both methods (Fig. 2B). DDA identified 3562 unique sites, whereas DIA identified 22568 unique sites (Fig. 2B). Both DDA and DIA workflows identified 3632 common ubiquitylated proteins, although both DDA and DIA also identified 460 and 3014 unique ubiquitylated proteins, respectively (Fig. 2D).

**Figure 2.**
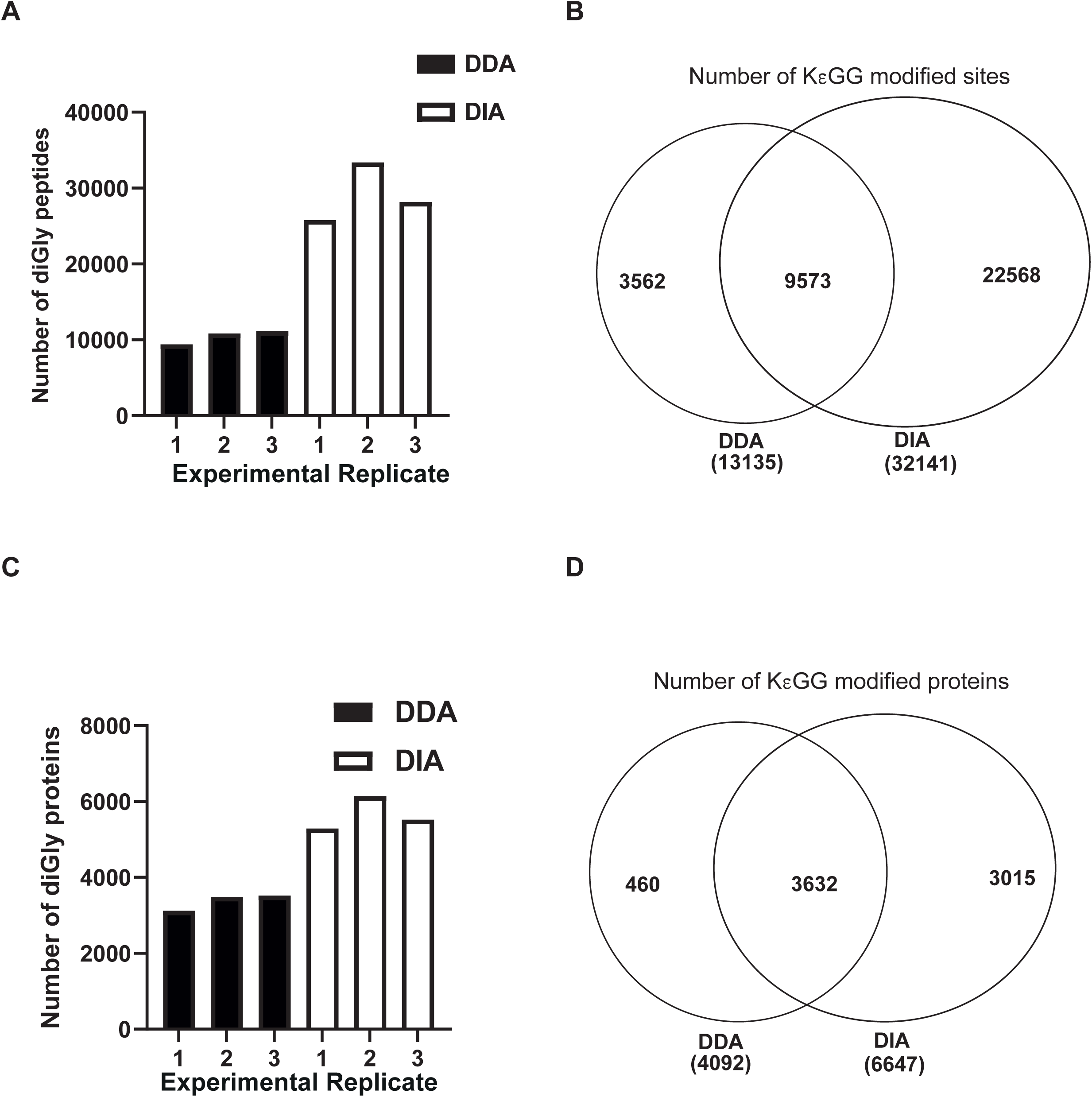
Comparison of data-dependant acquisition (DDA) vs. data-independent acquisition (DIA) for in-depth diGly ubiquitinome analysis. **(A)** Bar chart showing the number of diGly peptides identified from the 8M urea lysis buffer when analysed by DDA or DIA. The data shows the number of diGly peptides identified in each experimental replicate. **(B)** Venn diagram showing the total number of common and unique diGly modified sites identified using DDA or DIA methods. **(C)** As in (A), except the chart shows the numbers of diGly modified proteins identified in each experimental replicate when analysed by DDA or DIA. **(D)** As in (B), except the Venn diagram shows the diGly modified proteins.

### Comparing agarose vs. magnetic-bead diGly-enrichment methods for the identification of ubiquitylated peptides

Recent LC-MS/MS workflows allow the use of magnetic beads in automated settings for sample preparation and analysis. We therefore compared the performance of agarose beads vs. magnetic beads for diGly-peptide enrichment on cell extracts generated via urea-lysis method. Following enrichment from each method, the diGly peptides were analysed on Orbitrap Exploris 480 mass spectrometer using DIA method, and the DIA-NN search engine employed for data analysis. Three replicates from the agarose-bead diGly-enrichment identified 25775, 33400, 28195 diGly peptide spectra, corresponding to 5291, 6140, 5522 proteins, respectively (Fig. 3A&C; Supp. Data 3&4). In contrast, the three replicates from magnetic-bead diGly-enrichment yielded 33964, 36321 and 44622 diGly peptide spectra, corresponding to 5992, 6290 and 6901 proteins, respectively (Fig. 3A&C; Supp. Data 5). The magnetic-bead enrichment method outperforms the agarose-bead enrichment method in terms of the total number of diGly peptides (39865 vs. 32004) and ubiquitylated proteins (7414 vs. 6643) identified (Fig. 3B&D). From both methods, 28603 common diGly peptides corresponding to 6250 proteins were identified (Fig. 3B&D).

**Figure 3.**
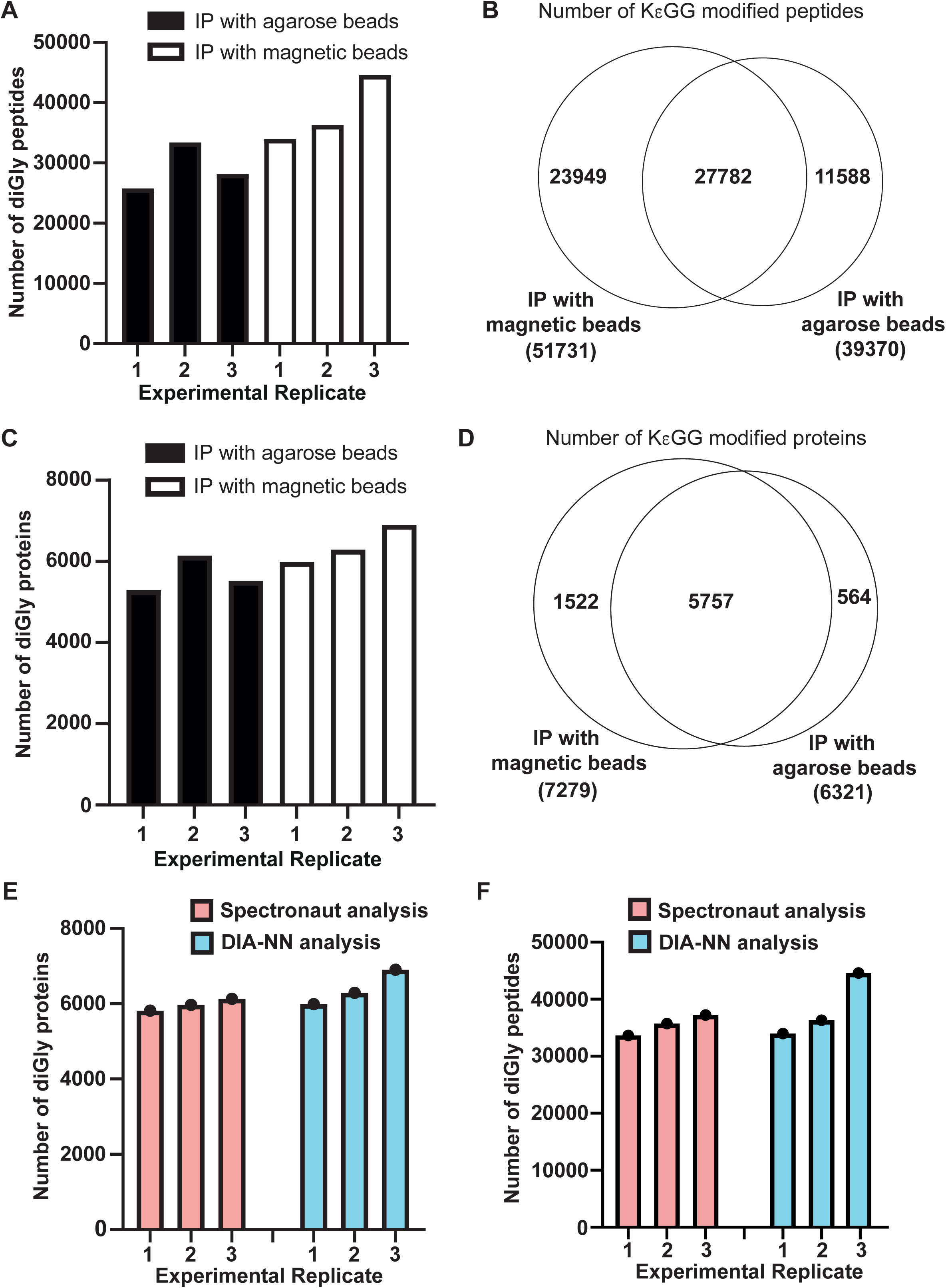
Comparison of agarose vs. magnetic bead enrichment of diGly peptides for ubiquitinome profiling by DIA. **(A)** Bar chart showing the number of diGly peptides identified by DIA-NN from each experimental replicate following enrichment with the anti-K-ε-GG antibody coupled to either agarose or magnetic beads. **(B)** Venn diagram showing the total number of common and unique diGly peptides identified in (A). **(C)** As in (A), except the chart shows the number of diGly modified proteins from each experimental replicate. **(D)** As in (B), except the Venn diagram shows diGly modified proteins identified in (C). **(E)** Bar chart showing the number of diGly modified peptides identified by Spectronaut or DIA-NN approaches from each experimental replicate following enrichment with the anti-K-ε-GG antibody coupled to the magnetic beads. **(F)** As in (E), except the chart shows the number of diGly modified proteins from each experimental replicate.

### Comparison of the Spectronaut™ Pulser and DIA-NN search engines

Using the data generated from the magnetic bead enrichment method above (Fig. 3A-D), we undertook a comparison of the two routinely used library-free search engines for DIA analysis, namely Spectronaut™ Pulser and DIA-NN. From three replicates, searches with Spectronaut™ Pulser identified 33625, 35663 and 37228 diGly peptide spectral matches, corresponding to 5819, 5966 and 6132 ubiquitylated proteins, respectively (Fig. 3E&F; Supp. Data 6). These were lower than the numbers identified by DIA-NN, which yielded 33964, 36321 and 44622 diGly peptide spectra, corresponding to 5992, 6290 and 6901 proteins, respectively (Fig. 3E&F; Supp. Data 5). Comparing these search engines, DIA-NN identified ∼21% more diGly precursors and 12.7% more ubiquitylated proteins than Spectronaut™ Pulser. Having established urea-lysis, magnetic-bead diGly enrichment and DIA-NN analysis as the optimal workflow for the most in-depth identification of ubiquitylated peptides and proteins from extracts, we set out to apply these methods to identify and map ubiquitylation changes that result from various TPD modalities.

### Mapping the ubiquitylation sites on the Tau-GFP induced by the VHL-aGFP16 AdPROM

Tau is a microtubule-associated protein that is primarily expressed in neurons of the central nervous system. Its main role is to stabilize microtubules and facilitate their assembly, which is essential for proper neuronal homeostasis. In Alzheimer’s disease (AD) and other tauopathies, however, Tau becomes hyperphosphorylated, leading to its detachment from microtubules and aggregation into neurofibrillary tangles (NFTs) ^36–38^. Here, we utilized the affinity-directed protein missile (AdPROM) system for targeted proteolysis ^39^ to target the degradation of Tau. For this, we used QBI-293 (cl. 4) cells stably expressing Tau40/P301L-GFP ^40^. In previous work, it was demonstrated that the fusion of VHL with an aGFP16 nanobody results in the recruitment of GFP-tagged proteins, such as VPS34 and PAWS1, to the CUL2-RBX1 E3 ligase complex for their targeted ubiquitylation, and subsequent degradation via the proteasome ^39^. Consequently, we hypothesized that in the presence of VHL-aGFP16 AdPROM, Tau40/P301L-GFP might also be recruited to the CUL2-RBX1 complex, leading to its ubiquitylation and degradation (Fig. 4A). Indeed, the introduction of VHL-aGFP16 AdPROM led to substantial reduction in levels of Tau40/P301L-GFP from QBI-293 (cl. 4) cells compared to control cells expressing the aGFP16 (Fig. 4B). Treatment of cells with the proteasome inhibitor MG132 robustly rescued the degradation of Tau40/P301L-GFP caused by VHL-aGFP16 (Fig. 4B). However, MG-132 also led to enhanced levels of Tau40/P301L-GFP in control cells expressing aGFP16, suggesting a naturally high turnover rate of Tau40/P301L-GFP in QBI-293 (cl. 4) cells (Fig. 4B). The Cullin neddylation inhibitor MLN4924 partially rescued the degradation of Tau40/P301L-GFP compared to DMSO control (Fig. 4B). A global total proteomic analysis using the DIA-NN workflow led to the identification of 8259 proteins from QBI-293 (cl. 4) cells expressing either VHL-aGFP16 AdPROM or aGFP16 control (Supp. Data 7, S1 A). A comparison of the proteome in the two cell types revealed that 34 proteins were significantly more and 22 less abundant in cells expressing the VHL-aGFP16 AdPROM compared to those expressing the aGFP16 control (Fig. 4C). Excitingly, there was a 7.5-fold reduction in levels of Tau in cells expressing the VHL-aGFP16 AdPROM compared to control cells, in all three independent replicates (Fig. 4C&D). As expected, we observed higher abundance of VHL in cells expressing VHL-aGFP16 AdPROM compared to control cells expressing aGFP16 control (Fig. 4C). Next, we sought to map the ubiquitylation sites on Tau40/P301L-GFP caused by the VHL-aGFP16 AdPROM using the optimised ubiquitylation proteomic workflow. For this, we performed global ubiquitylation analysis in MG-132 treated cells expressing either the VHL-aGFP16 AdPROM or the aGFP16 control. This analysis identified a total of 69,773 diGly peptide precursors, corresponding to 7733 ubiquitylated proteins (Fig. S1B & 4E, Supp. Data 8). Among the few ubiquitylated proteins that were identified to be upregulated in cells expressing VHL-aGFP16 AdPROM compared to those expressing aGFP16 control were Tau-GFP and VHL (Fig. 4E). Indeed, upon closer inspection, we identified 15 lysine residues on Tau and 12 lysine residues on GFP that showed enhanced ubiquitylation in cells expressing VHL-aGFP16 AdPROM compared to those expressing aGFP16 control (Fig. 4F, S1B&C.).

**Figure 4.**
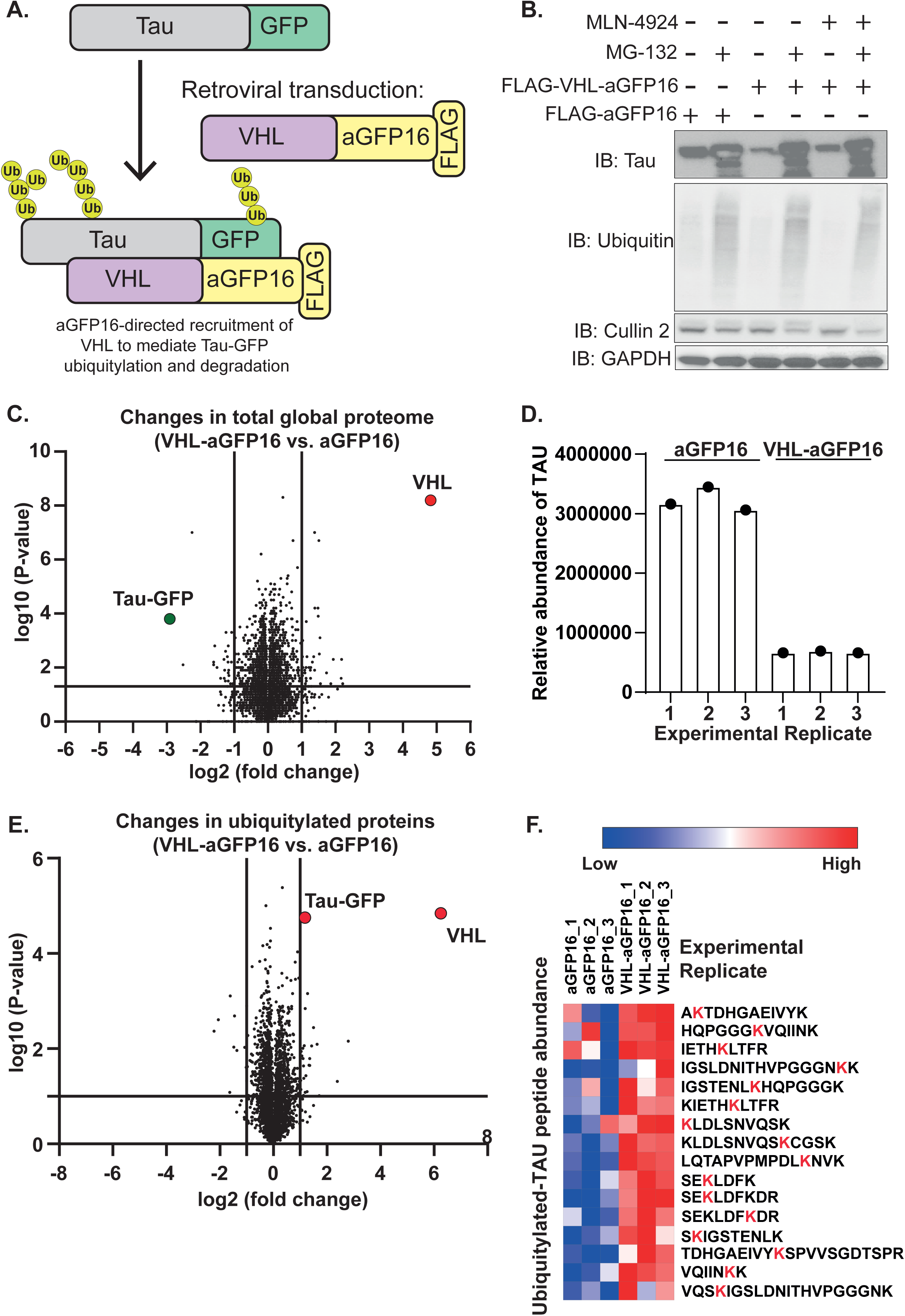
Identification of ubiquitylation sites on Tau-GFP caused by the VHL-aGFP16 AdPROM. (**A**) Schematic representation of the VHL-aGFP16 AdPROM-mediated ubiquitylation of Tau-GFP via proximity induction. (**B**) QBI-293 (cl. 4) cells stably expressing Tau-GFP were transduced with retroviruses encoding the VHL-aGFP16 or aGFP16 control AdPROMs as indicated. Cells were treated with DMSO (-), MG-132 (20 µM) or MLN-4924 (1 µM) for 6 h prior to lysis as indicated. Extracts (30 µg protein) were resolved by SDS-PAGE, and transferred to PVDF membranes, which were immunoblotted with the indicated antibodies. (**C**) Volcano plot showing a comprehensive total proteomic analysis from extracts of cells expressing VHL-aGFP16 AdPROM compared to those expressing the aGFP16 control. Tau-GFP and VHL are indicated. (**D**) Quantitative analysis of Tau protein levels derived from mass spectrometry analysis from across individual replicates of QBI-293 (cl. 4) cells stably expressing Tau-GFP transduced with retroviruses encoding the VHL-aGFP16 or aGFP16 AdPROMs as indicated. (**E**) Volcano plot showing global ubiquitylated proteome changes in extracts of cells expressing VHL-aGFP16 compared to those expressing the aGFP16 control, in the presence of the proteasomal inhibitor MG-132. Tau-GFP and VHL were both identified as ubiquitylated proteins in cells expressing VHL-aGFP16 AdPROM. (**F**) Heatmap of the ubiquitylated lysine residues identified on Tau-GFP in QBI-293 (cl. 4) cells transduced with retroviruses encoding the VHL-aGFP16 or aGFP16 AdPROMs as indicated in the presence of the proteasome inhibitor MG-132. The colour spectrum bar shows relative abundance.

### Mapping ubiquitin sites on endogenously knocked-in dTAG-PPP2CA induced by the PROTAC dTAG-13

Next, to apply the optimised proteomics workflow for TPD at the endogenous level, we sought to investigate the alterations in global ubiquitylation upon degradation of dTAG-PPP2CA and potentially map direct ubiquitylation sites on dTAG-PPP2CA in *^dTAG/dTAG^PPP2CA* knock-in (KI) HEK293 cells induced by the PROTAC dTAG-13 ^30^ (Fig. 5A). PPP2CA is the principle catalytic subunit of the protein serine-threonine phosphatase 2A (PP2A) holoenzyme complex, which plays a crucial role in a plethora of cellular processes, encompassing cell growth, differentiation, cell cycle progression, and apoptosis ^41^. Previously, we showed that the selective degradation of dTAG-PPP2CA by dTAG-13 in *^dTAG/dTAG^PPP2CA* knock-in HEK293 cells identified >2000 putative protein substrates of PPP2CA ^30^. Treatment of *^dTAG/dTAG^PPP2CA* KI HEK293 cells with 100 nM dTAG-13 for 16 h led to a robust degradation of dTAG-PPP2CA compared to DMSO treated controls, while this degradation was partially rescued by the proteasomal inhibitor MG-132 (Fig. 5B). A global proteomic analysis of DMSO and dTAG-13-treated cells in the absence of MG-132 treatment using DIA-NN identified a total of 10,207 proteins (Supp. Data 9). Remarkably, PPP2CA and two members of the PP2A holoenzyme complexes, namely PPP2R5D and PPP2R5A, were selectively and significantly downregulated by dTAG-13 over DMSO control treatment (Fig. 5C). We detected at least 4-fold lower abundance of PPP2CA upon dTAG-13 treatment compared to DMSO in all 3 replicates (Fig. 5D). Next, we carried out global ubiquitinome analysis in these samples. A total of 43,416 ubiquitylated precursors corresponding to 6680 proteins were identified (Fig. 5E; Supp. Data 10). Although both western blot and total proteomic data showed a near-complete degradation of dTAG-PPP2CA by dTAG-13 (Fig. 5B&D), a total of 7 ubiquitylated sites were identified on PPP2CA upon dTAG-13 treatment (Fig. S2A). Analysis of the global ubiquitylated peptides showed that 972 ubiquitylated peptides corresponding to 502 proteins were significantly more abundant and 962 peptides corresponding to 780 proteins were less abundant upon dTAG-PPP2CA degradation by dTAG-13 compared to DMSO controls (Fig. 5E&F, S2B). This was quite surprising, given the total proteomic analysis showed very few changes in abundance (Fig. 5C), suggesting that these alterations protein ubiquitylation caused by degradation of PPP2CA do not confer degradation signals. Nonetheless, further bioinformatic analysis of proteins with enhanced ubiquitylation showed distribution in many subcellular compartments (Fig. S2C) and involvement in many biological processes and molecular functions (Fig. S2D-J). From the seven ubiquitylated sites identified on PPP2CA, only three were unique to PPP2CA, while the remaining four were also conserved on PPP2CB. To determine the ubiquitylation stoichiometry, we calculated the ratios of 3 unique diGly-modified peptides mapped to K8, K21 and K29 of PPP2CA which showed significant increase in abundance upon dTAG-13 treatment compared to DMSO controls relative to protein abundance in each experimental replicate (Fig. 5G-I). This analysis revealed a very high level of ubiquitylation at these sites upon dTAG-13 treatment compared to DMSO controls in all 3 replicates (Fig. 5G-I). We also acquired proteome and ubiquitinome data from cells were treated with DMSO or dTAG-13 in the presence of MG-132 (Fig. 5B). The proteomics analysis of these cells identified a total of 10,579 proteins with few significant differences between DMSO and dTAG-13 treatment conditions (Fig. S3A, Supp. Data 11). The global ubiquitylation analysis of the MG-132 treated cells were identified 73,974 DiGly peptides corresponding to 8,115 proteins (Fig. S3 B, Supp. Data 12). A total of 8 ubiquitylated peptides on PPP2CA were mapped, including those ubiquitylated on K8, K21 and K29, were significantly upregulated in MG-132 treated extracts upon dTAG-13 treatment compared to DMSO treated controls (Fig. S3B). The fold changes in the abundance of the identified PPP2CA ubiquitylated peptides in dTAG-13 treated cells compared to treated DMSO controls were more pronounced in the absence of MG-132 treatment, suggesting that MG-132 treatment promotes the accumulation of ubiquitylation on dTAG-PPP2CA potentially caused due to its natural turnover rate.

**Figure 5.**
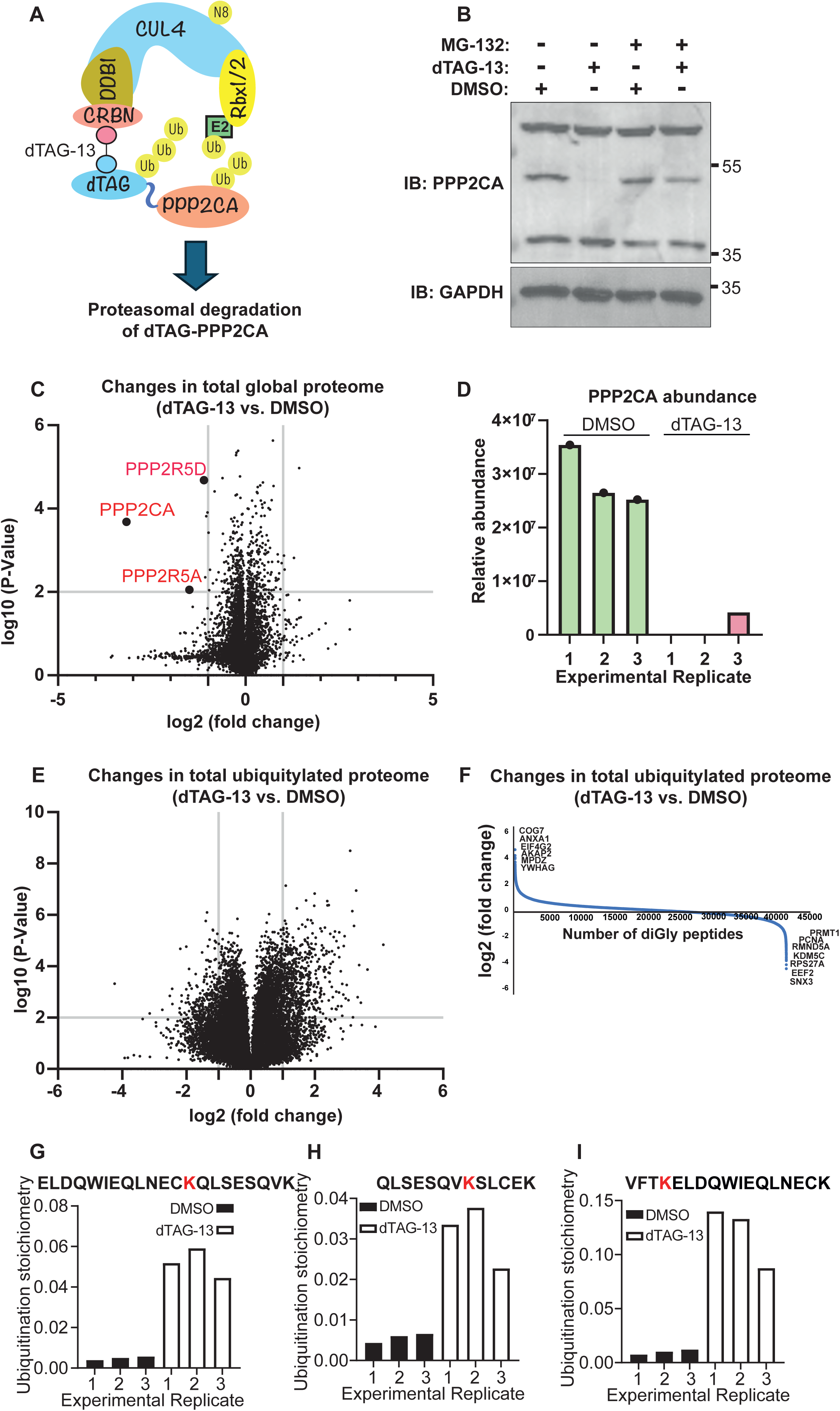
Identification of ubiquitylation sites on dTAG-PPP2CA at the endogenous level induced by the PROTAC dTAG-13. **(A)** Cartoon of the PROTAC dTAG-13 depicting how it recruits the CUL4A^CRBN^ E3 ligase complex to ubiquitylate dTAG-PPP2CA for its degradation through the ubiquitin-proteasome pathway. **(B)** Homozygous *^dTAG/dTAG^PPP2CA* knock-in HEK293 cells were treated with DMSO, dTAG-13 (100 nM) or MG-132 (20 µM) as indicated for 16 h prior to lysis. Extracts (30 µg protein) were resolved by SDS-PAGE, and transferred to PVDF membranes, which were immunoblotted with the indicated antibodies. **(C)** Volcano plot showing changes in abundance of proteins identified in *^dTAG/dTAG^PPP2CA* knock-in HEK293 cells treated with dTAG-13 compared to DMSO control. Indicated are components of the PP2A holoenzyme complexes that are degraded by dTAG-13 significantly compared to DMSO control. **(D)** A quantitative analysis of PPP2CA protein levels from *^dTAG/dTAG^PPP2CA* knock-in HEK293 cells treated with dTAG-13 or DMSO from across individual replicates as indicated. **(E)** Volcano plot showing changes in ubiquitylated proteins identified from *^dTAG/dTAG^PPP2CA* knock-in HEK293 cells treated with dTAG-13 compared to DMSO in the absence of MG-132 treatment. **(F)** As in (E) except that the data is represented as a trend plot. Ubiquitylated proteins with most differences (lower or higher) in abundance are indicated. **(G-I)** The abundance of PPP2CA-K21 (G), PPP2CA-K29 (H), and PPP2CA-K8 (I) ubiquitylation in *^dTAG/dTAG^PPP2CA* knock-in HEK293 cells treated with dTAG-13 compared to DMSO from individual replicates.

To capture early ubiquitylation events on dTAG-PPP2CA upon dTAG-13 treatment when degradation begins to be detected, we performed a shorter time course of dTAG-13 treatment for 15 min, 30 min and 60 min prior to lysis and analysed global ubiquitylation with DIA-NN (Fig. 6A-C). In total, we identified six lysine residues on PPP2CA which were significantly ubiquitylated upon dTAG-13 treatment compared to DMSO controls (Fig. 6A-C, Supp. Data 13). While 3 of these peptides are unique to PPP2CA, 3 are also conserved with PPP2CB, although dTAG-13 did not induce any degradation of endogenous PPP2CB (Fig. 5C; Supp. Data 13). For each ubiquitylation site, we measured abundance across each dTAG-13 treatment time course relative to DMSO control (Fig. 6D). The ubiquitylation on K8 showed a time-dependent increase in abundance from 15 min to 60 min following dTAG-13 treatment (Fig. 6D). For all other peptides, ubiquitylation levels peaked at 15 min and reduced in abundance by 60 min following dTAG-13 treatment (Fig. 6D; Supp. Data 13). When all the ubiquitylated residues were mapped to PPP2CA within the structure of the PP2A holoenzyme complex ^42^(PDB: 7CUN), all sites reside in a common region (Fig. 6 E), supporting the notion that dTAG-13 induces the recruitment of the CUL4A^CRBN^ E3 ligase complex to this region.

**Figure 6.**
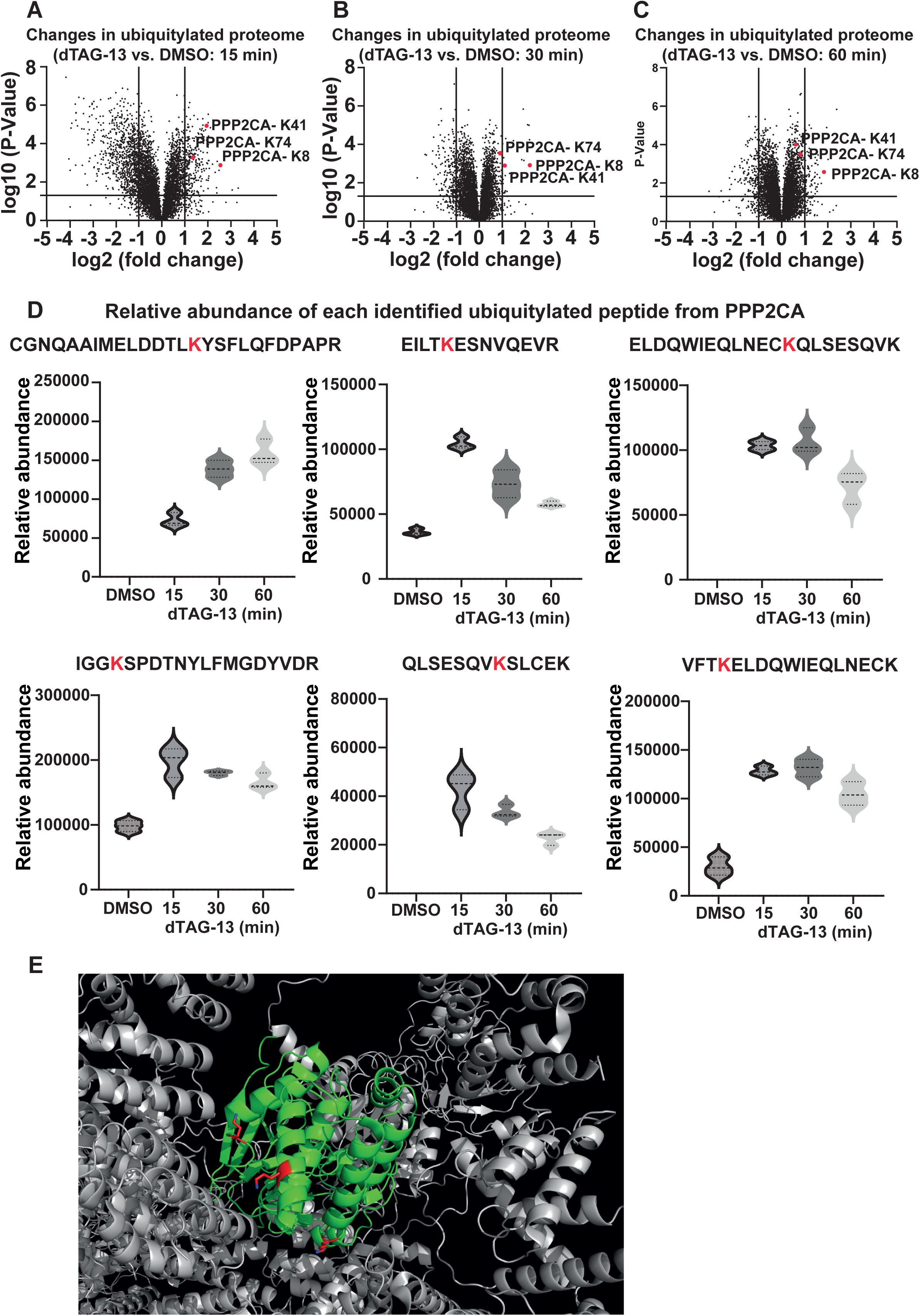
Identification of ubiquitylation sites on dTAG-PPP2CA at the endogenous level induced by a short treatment of dTAG-13 PROTAC. **(A)** Volcano plot representing changes in global ubiquitylated sites in *^dTAG/dTAG^PPP2CA* knock-in HEK293 cells treated for 15 min with dTAG-13 compared to DMSO controls. Sites identified on PPP2CA upon dTAG-13 treatment are indicated. **(B)** As in (A) except that dTAG-13 was applied for 30 min. **(C)** As in (A) except that dTAG-13 was applied for 60 min. **(D)** The changes in abundance of the indicated ubiquitylated peptides that were identified on PPP2CA after 15 min, 30 min and 60 min of dTAG-13 treatment compared to DMSO treatment. **(E)** The 3D-structure of the PPP2CA rendered from the Integrator-PP2A complex (PDB: 7CUN) with annotations of dTAG-13-induced ubiquitylation sites on PPP2CA in red.

### Mapping the ubiquitylated sites on endogenous SGK3 induced by SGK3-PROTAC1

Next, we sought to employ the optimised proteomic workflow to potentially map PROTAC-induced ubiquitylation sites on an endogenous protein substrate. We chose the SGK3-PROTAC1, which was recently reported to cause an efficient and selective degradation of endogenous SGK3 in cells ^31^. Treatment of HEK293 cells with 100 nM SGK3-PROTAC1 for 8 h caused almost complete degradation of SGK-3 compared to DMSO control (Fig. 7A). Treatment of cells with MG-132 partially rescued the degradation of SGK-3 by SGK3-PROTAC1 (Fig. 7A). Total proteomic analysis of SGK3-PROTAC1 treated cells compared to DMSO control using DIA showed a robust and significant degradation of SGK3 (Fig. 7B; Supp. Data 14). We also carried out diGly enrichment and ubiquitinome analysis by DIA on SGK3-PROTAC1 and DMSO treated samples. A total of 17,636 ubiquitylated precursors corresponding to 3906 proteins were identified (Fig. S4A; Supp. Data 15). In these samples, we were unable to identify any ubiquitylated peptides corresponding to SGK3, potentially corroborating the fact that SGK3 after 8 h treatment with SGK3-PROTAC1 was completely degraded. To preserve PROTAC-induced ubiquitylated SGK3 from proteasomal degradation, we employed the proteasomal inhibitor MG-132. Proteomic analysis of these samples identified 7779 number of proteins (Fig. S4 B Supp. Data 16). For ubiquitylation proteomic analysis using DIA, we compared SGK3-PROTAC1+MG-132-treated samples against DMSO+MG-132-treated controls. In these extracts, we identified a total of 42791 ubiquitylated peptide precursors corresponding to 6279 proteins (Fig. 7C&D; Supp. Data 17). We identified and quantifiably mapped 11 ubiquitylated lysine residues on SGK3. From these, 8 ubiquitylated peptides were significantly higher in abundance in SGK3-PROTAC1+MG-132-treated samples compared to DMSO+MG-132-treated controls (Fig. 7E). As the SGK3-PROTAC1 binds SGK3 on its ATP-binding pocket and recruits CUL2^VHL^ E3 ligase complex, it was perhaps not surprising that all 8 ubiquitylated lysine residues mapped to the kinase domain of SGK3, as denoted on the structure of SGK3 (Fig. 7F; PDB: 6EDX). These data demonstrate that our ubiquitinome proteomics workflow can identify the PROTAC-modified lysine residues on endogenous target proteins.

**Figure 7.**
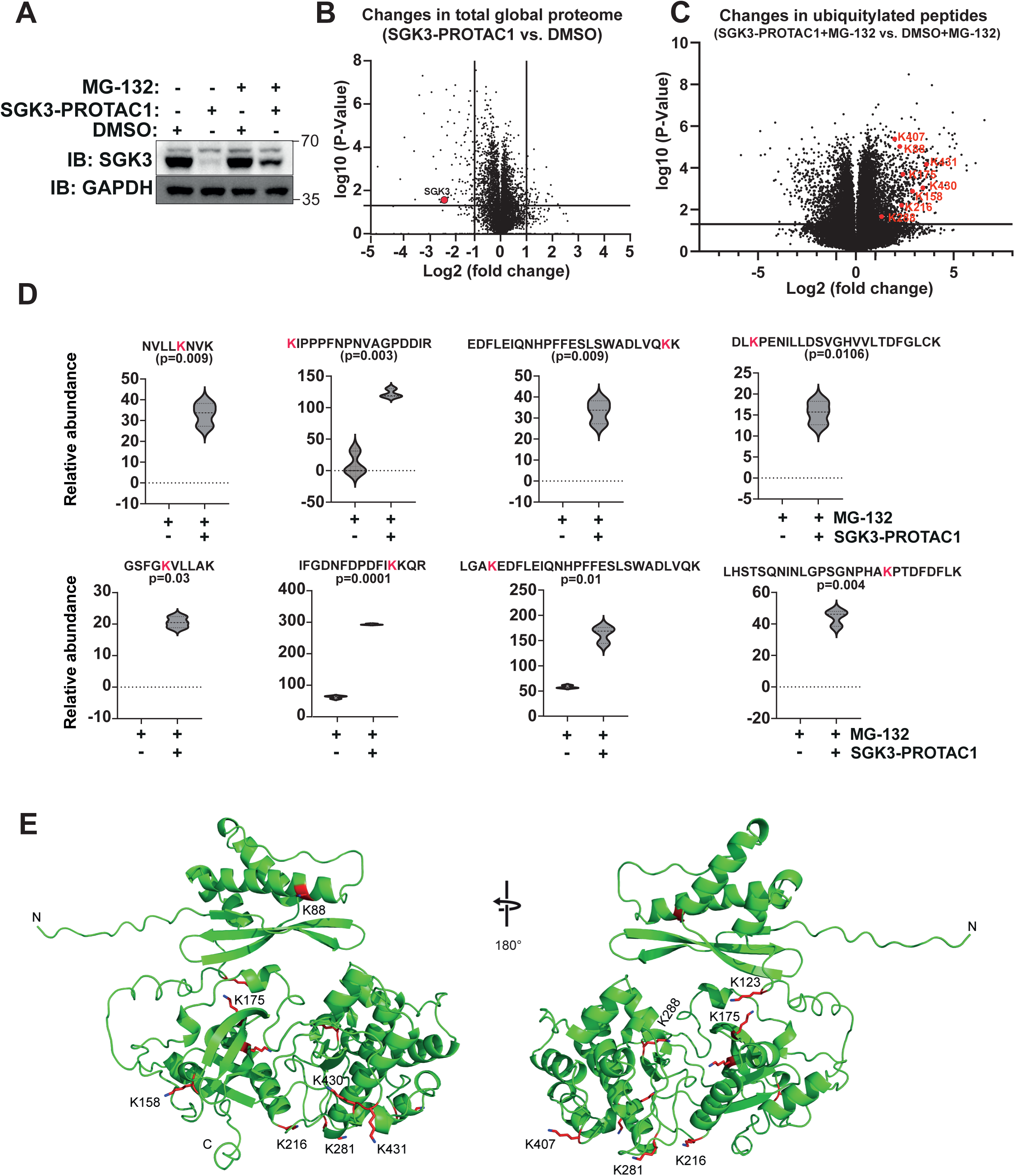
Identification of ubiquitylation sites on SGK3 induced by SGK-3 PROTAC1. **(A)** HEK293 cells were treated with DMSO, SGK3-PROTAC1 (100 nM) or MG-132 (20 µM) as indicated for 16 h prior to lysis. Extracts (30 µg protein) were resolved by SDS-PAGE, and transferred to PVDF membranes, which were immunoblotted with the indicated antibodies. **(B)** Volcano plot showing global changes in the proteome abundance from HEK293 cells treated with SGK3-PROTAC1 compared to DMSO control in the absence of MG-132. **(C)** Volcano plot representing global changes in the abundance of ubiquitylated peptides from HEK293 cells treated with SGK3-PROTAC1 compared to DMSO control in presence of MG-132. The ubiquitylated SGK3 peptides that were identified and show significant increase upon SGK3-PROTAC1 treatment are indicated. **(D)** The trends in abundance of all identified ubiquitylated peptides from SGK3 from HEK293 cells treated with SGK3-PROTAC1+MG-132 compared to DMSO+MG-132 controls are indicated. **(E)** All ubiquitylation sites on SGK3 identified in (D) shown on the 3D-structure of SGK3 (PDB: 6EDX).

## Discussion

TPD has emerged as a highly promising therapeutic modality for many human diseases, with several clinical trials ongoing ^43,44^. The ability to selectively eliminate disease-causing proteins by hijacking the cellular ubiquitin-proteasome system offers new avenues for drug development against targets that were until recently considered intractable. Many TPD modalities, including AdPROM, PROTACs and molecular glues, seek to induce proximity between the target protein and an E3 ligase to facilitate the polyubiquitylation and degradation of the target protein. Therefore, it is important to understand precisely how degraders induce ubiquitylation of target proteins, and whether redirecting an E3 ligase to new targets affects the turnover of its natural substrates.

In recent years, there have been significant advances in proteomic workflows that have improved both throughput and depth of coverage ^45–47^. For ubiquitinome proteomics, the ability to enrich diGly remnant peptides transformed our ability to identify ubiquitylated peptides by mass spectrometry. Several workflows have been reported for ubiquitinome analysis, with variable outcomes. In this study, we compared several state-of-the-art methodologies for ubiquitinome proteomics and report an optimized workflow. Based on our side-by-side comparisons, we recommend employing a ubiquitinome proteomic workflow based on urea-based lysis, enrichment of diGly peptides using magnetic beads, and analysis of the mass-spectrometry data using DIA-NN for optimal depth of coverage. However, while the majority of ubiquitylated peptides and proteins identified by each of the urea or SDS lysis, DIA or DDA analysis and agarose or magnetic bead enrichment methods were common, there were still unique ubiquitylated peptides and proteins identified by each method, even for those where the depth of coverage was lower. Although some of the differences are likely to be due to technical inconsistencies and differential protein ubiquitylation stoichiometries across biological replicates, analysing the reproducible coverage of unique ubiquitylated peptides and proteins identified by each methodology, especially those yielding lower overall coverage, might still help researchers choose specific methodologies that turn out to be more suitable for detecting specific ubiquitylated peptides or proteins.

We employed the optimised ubiquitinome workflow to identify ubiquitylation sites on substrate proteins and global changes in ubiquitylation upon application of different TPD modalities. The VHL-aGFP16 AdPROM-mediated targeted degradation of Tau-GFP identified 15 lysine residues on TAU and 12 lysine residues on GFP that were ubiquitylated, while there was minimal change in global ubiquitylation. For this, we had to employ the proteasome inhibitor MG-132 to preserve ubiquitylated Tau-GFP caused by VHL-aGFP16 AdPROM prior to its destruction by the proteasome. dTAG-13-induced degradation of endogenously knocked-in dTAG-PPP2CA also identified multiple ubiquitylated lysine residues on PPP2CA, even in the absence of MG-132 treatment. Here, we also explored and documented the dynamic changes in peptide level ubiquitylation on 3 different PPP2CA residues over a short course of dTAG-13 treatment for up to 60 minutes. Finally, we were able to determine multiple ubiquitylation sites on endogenous SGK3 induced by SGK3-PROTAC1. However, for SGK3, as with AdPROM-mediated Tau-GFP degradation, we needed to prevent the degradation of ubiquitylated SGK3 by employing the proteasomal inhibitor MG-132 in order to enrich and identify ubiquitylated peptides on SGK3. As MG-132 inhibits the proteasome and results in the accumulation of ubiquitylated proteins globally, it may trigger accumulation of ubiquitylation on the same residues of target proteins with high natural turnover rates as those that degraders would otherwise trigger. In such cases, MG-132 treatment can compound identification of ubiquitylation events triggered by the degrader. To overcome the potential issues with MG-132 enrichment, our data obtained following short-term treatment of cells harbouring dTAG-PPP2CA knockin with dTAG-13 in the absence of MG-132 treatment demonstrates that ubiquitylation sites on target proteins can be mapped following a short treatment course with PROTACs. Eventually, whether to employ MG-132 for enrichment of PROTAC-induced ubiquitylated proteome could be informed by degradation kinetics and natural turnover rates of the target protein.

One recurring theme we observed when analysing global changes in the abundance of ubiquitylated peptides and total protein by degraders was that there were thousands of ubiquitylated peptides that significantly altered in abundance upon degrader treatment. Surprisingly, this did not translate into significant changes in protein abundance, as for most degraders, degradation of the target protein was mostly selective. It is possible that altered ubiquitylation observed is a consequence of target protein degradation but that the stoichiometry of ubiquitylation on individual target proteins is too low to cause substantial changes in total protein abundance. Another possibility is that, in addition to K48-linked chains mediating degradation, other types of ubiquitin linkages may be involved, contributing to cellular signaling rather than proteasomal degradation. Similarly, some altered ubiquitylation events could be due to aberrant E3 ligase activity from overexpression of VHL-aGFP16 (for AdPROM-mediated degradation of Tau-GFP) or after redirecting CRBN (dTAG-13) and VHL (SGK3-PROTAC1) to target the degradation of dTAG-PPP2CA and SGK-3, respectively. More unbiased global ubiquitylation data on new generations of PROTACs may shed new light on this in the future.

The ability to rapidly establish the mode of action for PROTACs and molecular glues with regard to target protein degradation is of great significance, especially to glean mechanistic insights and to potentially predict modes of degrader-resistance. By identifying the specific ubiquitylation sites on target proteins and providing the atlas of ubiquitylation changes, our workflow enables the characterization of key ubiquitylation events that govern the degradation of disease-associated proteins and offers clues to the potential global consequences of either redirecting the E3 ligase or degradation of the target protein. This knowledge is instrumental in rational design and optimization of targeted protein degradation strategies. By precisely and efficiently profiling ubiquitin sites on target proteins, our approach facilitates a comprehensive understanding of the mechanisms underlying how small molecular degraders, such as PROTACs (proteolysis-targeting chimeras) and molecular glues, work.

## Materials and Methods

### Cell culture, manipulation, and lysis

The cell lines used in this study were: human embryo kidney HEK293 cells (ATCC, CRL-1573), HEK293-FT cells (Invitrogen, Cat# R70007), HEK293-derived QBI-293 (Clone 4.0) cell line stably expressing full length human Tau (T40-2N4R) carrying the P301L mutation with a GFP-tag at the C-terminus ^40^ and *^dTAG/dTAG^PPP2CA* knock-in HEK293 cells generated using CRISPR/Cas9 genome editing ^30^. All cells were cultured in DMEM (Life Technologies) supplemented with 10% (v/v) foetal bovine serum (FBS, Thermo Fisher Scientific), 2 mM L-glutamine (Lonza), 100 U/mL penicillin (Lonza) and 0.1 mg/mL streptomycin (Lonza). Cells were maintained at 37°C with 5% CO_2_ in a water-saturated incubator and regularly tested for mycoplasma contamination. For passaging, trypsin/EDTA was used at 37°C to detach cells. Cells were treated with either DMSO control or specific concentrations of the indicated compounds (which were diluted in DMSO) for the indicated times, washed twice with ice-cold PBS, and lysed by scrapping in freshly prepared sodium dodecyl sulphate (SDS) lysis buffer or urea lysis buffer. The SDS lysis buffer contained 2% SDS, 10 mM (TCEP-C4706), 40 mM CAA (22790), 50 mM TEABC at pH8.5. The urea lysis buffer contained 8 M urea in 50 mM TEABC. Both lysis buffers contained protease and phosphatase inhibitor cocktail tablets. The SDS lysates were heated to 95 °C for 10 min and then sonicated using an ultrasonic probe device (1 min, energy output ∼40%). The urea lysis and subsequent steps were done at room temperature to avoid carbamylation. The urea lysates were also sonicated. Both sets of lysates were centrifuged at 12,000 rpm for 10 minutes, and supernatants were used for the in-solution digestion, as detailed in Fig. 1. For proteasome inhibition, the respective cells were treated with 20 µM MG-132 at ∼80% confluence for 6 h prior to lysis. The protein concentration was measured using the BCA protein estimation assay.

### Retroviral generation of stable cell lines

Retroviral AdPROM constructs were generated using pBABED-puro plasmids, as described previously ^39^. To produce AdPROM retrovirus particles, 6 μg of the plasmid was transiently transfected into a 10-cm dish of approximately 70% confluent human embryonic kidney (HEK) 293FT cells, along with 2.2 μg of pCMV5-VSV-G and 3.8 μg of pCMV5-GAG/POL. For transfection, plasmids were first mixed in 300 μl of Opti-MEM in one tube. In a separate tube, 24 μl of 1 mg/ml PEI was added to 300 μl of Opti-MEM. Both mixtures were incubated for 5 minutes before combining and, after brief mixing, they were left to incubate for an additional 20 minutes. The resulting transfection mixture was then added to 9 ml of complete DMEM and applied to the HEK293FT cells. After 16 h, the transfection medium was replaced with fresh culture medium. Viral supernatant was collected 24 h later, filtered through a 0.45 μm sterile syringe filter, and used for transduction. Target cells at approximately 60% confluency were transduced using the optimized titre of retroviral medium diluted in fresh medium containing 8 μg/ml polybrene (Sigma-Aldrich) for 24 h. Following transduction, the medium was replaced with selection medium containing 2 μg/ml puromycin to enrich cells carrying the retroviral particles. After complete elimination of non-transduced cells under selection, the surviving pool of transduced cells was used for subsequent experiments.

### Immunoblotting

Reduced and denatured cell lysates containing equal amounts of protein (30 μg) were resolved by SDS-PAGE and transferred to PVDF membranes. After blocking the membranes in 5% (w/v) non-fat milk in TBS-T (20 mM Tris, 150 mM NaCl, 0.1% Tween-20) for 1 h at room temperature, membranes were incubated with primary antibody diluted in 5% milk/TBS-T overnight at 4°C in a shaker. Membranes were then washed 3×10 min in TBS-T with constant shaking and subsequently incubated with HRP-conjugated secondary antibody diluted in 5% milk/TBS-T solution for 1 h at room temperature. Membranes were then washed 3×10 min in TBS-T and the signal detection via chemiluminescence (Promega) was performed using ChemiDoc imaging system (Bio-Rad).

### Crosslinking of K-ε-GG antibody

The cross-linking of the anti-K-ε-GG antibody to agarose beads was carried out as described previously ^15^. The anti-K-ε-GG antibody was purchased as part of the PTMScan® ubiquitin remnant motif (K-ε-GG) kit from Cell Signaling Technology (#5562). Briefly, the entire vial of anti-K-ε-GG antibody coupled to agarose beads were washed three times with 100 mm sodium borate, pH9.0 in 1.5 ml Eppendorf tubes. After washing, cross-linking was performed by resuspending anti-K-ε-GG-beads in 1 ml of 20 mM dimethyl pimelimidate (DMP) and incubated at room temperature for 30 min on a rotating platform. To stop the reaction, beads were washed twice with 1 ml of 200 mM ethanolamine (pH8.0) and subsequently incubated in 1 ml of 200 mM ethanolamine for 2 h at 4°C with rotation. After blocking, the anti-K-ε-GG-beads were washed three times in 1 ml of ice-cold immunoprecipitation buffer (IP) (50 mm MOPS, pH 7.2, 10 mm sodium phosphate, 50 mm NaCl), resuspended in IP buffer, and stored at 4°C for future use.

### In solution digestion and peptide clean-up

After urea/SDS lysis, the protein content in the extracts was measured by the BCA protein estimation method. The extracts with indicated amount of protein were subjected to an in-solution digestion. For SDS lysates, the S-Trap™ midi-spin column digestion protocol was used ^33^. Briefly, for clarified SDS extracts, the SDS concentration was adjusted to 5% by using 10% SDS stock. Extracts were reduced in 5 mM DTT for 60 min at room temperature and alkylated using 10 mM chloroacetamide (CAA) for 20 min at room temperature in the dark. 1/10 volume of 12% phosphoric acid was added to the extracts, followed by 7-fold volume of wash-buffer containing (90% methanol in 10% 1M TEABC). This final extract was passed through the S-Trap™ column. The columns were washed 4 times with 3 ml of the wash buffer. 1:10 ratio of Trypsin:extract protein (by mass) and 1:100 ratio of LysC:extract protein (by mass) was employed for in-column digestion. The column was incubated at 47 °C for 1 h, followed by 12 h at room temperature. The digested peptides were eluted using 50 mM TEABC, 0.1% formic acid followed by 50% ACN. The eluate was dried using SpeedVac and reconstituted in solution for diGly enrichment. For urea lysis, indicated amounts of extract proteins were reduced in 5 mM DTT for 60 min at room temperature and alkylated using 10 mM CAA for 20 min at room temperature in the dark. The extracts were then incubated with LysC for 4 h at room temperature at 1:100 ratio of enzyme:extract protein (by mass). Following LysC digestion, the lysates were diluted with 50 mM TEABC to reduce the concentration of urea to less than 2 M. Trypsin at 1:20 ratio of enzyme:extract protein (by mass) was used for the in-solution digestion and incubated for 16 h at room temperature. The reaction was quenched with the addition of trifluoroacetic acid (TFA) to give a final concentration of 1% TFA (v/v) and samples were desalted on 200 mg Sep-Pak C_18_ cartridges (Waters). For Sep-Pak clean-up, the following solvents were prepared fresh: activation solvent (100% (v/v) acetonitrile (ACN)); 5 mL Solvent-1 (0.1% (v/v) TFA in ACN); 5 mL Solvent-2 (0.1% (v/v) formic acid (FA) in ACN); 5 mL Solvent-3 (50% (v/v) ACN in 0.1% (v/v) FA). Sep-Pak cartridges were equilibrated with 5 mL activation solvent (100% CAN), followed by 5 mL Solvent-1, and finally with 5 mL of Solvent-2 twice. Samples were then loaded onto the equilibrated C_18_ cartridges, washed with 5 mL 0.1% TFA four times, followed by washing with 5 mL 0.1% FA. Samples were then eluted with 6 mL 50% ACN, 0.1% FA. Desalted samples were then dried completely in a SpeedVac concentrator.

### diGly peptide enrichment

The agarose beads cross-linked to anti-K-ε-GG antibody were equilibrated in PTMScan® HS Immunoaffinity Purification (IAP) Wash/Bind Buffer (CST #42424). The dried digested peptides were dissolved in 1 mL of IAP buffer, centrifuged at 12,000 rpm for 10 min, and the supernatant was added to the equilibrated agarose beads cross-linked to anti-K-ε-GG antibody (one-eighth of the volume of the cross-linked antibody-agarose bead slurry). The IP mix was rotated at 4°C for 2 h with end-over-end rotation. The agarose beads were then washed twice with 1 mL of IAP buffer, followed by an additional wash with 1 mL of ice-cold Milli-Q® water. After removing all the supernatant, the beads were incubated with 200 µL of 0.15% TFA at room temperature while shaking at 1,400 rpm. After a brief spin, the supernatant was recovered and desalted using freshly prepared 200 µL two-plug StageTips. The eluted peptides were dried in SpeedVac and stored in −80°C until mass spectrometry analysis. For Magnetic beads conjugated to the anti-K-ε-GG antibody (PTMScan® HS Ubiquitin/SUMO Remnant Motif Kit #59322) were washed 3 times in 1 mL ice-cold PBS and reconstituted in 200 μL of PBS. 5 μL of magnetic beads were directly added to the peptide supernatant, obtained from dried digested peptides which were reconstituted in 1 mL of IAP buffer by repeat pipetting and clarified by centrifugation at 10,000xg for 5 min at 4°C. The magnetic anti-K-ε-GG antibody bead-peptide mix was incubated on an end-over-end rotating platform for 2 h at 4°C. The tube was then placed in a magnetic stand for 10 seconds, and the unbound peptide solution was discarded. The magnetic beads were washed with 1 mL of IAP buffer 3 times followed by 2 washes with 1 mL ice-cold Milli-Q® water. The beads were then incubated with 0.15% TFA and the diGly modified peptides were collected. The peptide solution was subjected to the stage-tip based C_18_ clean-up. The peptides were reconstituted in 0.1% formic acid. The peptides were passed through C_18_ stage tip that were activated by passing 30 μl of 100% ACN and equilibrated with 0.1% formic acid. The column was washed again 2 times with 0.1% formic acid and the peptides were eluted in an Eppendorf tube by using 50% ACN/0.1% formic acid. The eluted peptides were dried in SpeedVac and stored in −80°C until mass spectrometry analysis.

### LC-MS/MS analysis

The eluted peptides were analysed on an Orbitrap Exploris 480 mass spectrometer interfaced with Dionex Ultimate 3000 nanoflow liquid chromatography system. The dried peptides were reconstituted in 0.1% formic acid and loaded on a trap column (Acclaim PepMap100 C_18_, 100 μm × 2 cm, 5 μm, 100 Å) and separated on an analytical column (Acclaim PepMap RSLC C_18_, 75 μm × 50 cm, 2 μm, 100 Å) at a flow rate of 250 nL/min using a step gradient of 0-8% solvent B (90% ACN/0.1% FA) for the first 12 min, followed by 8-25% up to 110 min, and 25-37% for 110-125 min. The total run time was set to 145 min. For DDA analysis MS scan resolution was set at 120,000, and normalised AGC target was 300%. For MS2, the ions were fragmented using HCD with collision energy of 30, AGC target was set at 3000%, and resolution was set at 30,000.

For DIA MS, scan resolution was set at 120,000, scan range was set between 375-1500, and normalised AGC target was set at 300%. For MS2 the ions were fragmented using HCD with stepped energy of 10 and 25, AGC target set at 3000%, and variable windows were used for MS2 acquisition. These windows are provided in Supplementary Information.

### Raw data processing

The DDA proteomic raw data were searched using SEQUEST HT search engines with Proteome Discoverer 2.4 (Thermo Fisher Scientific). The following parameters were used for searches: Database: human proteome from Uniport, Precursor mass tolerance 10 ppm, Fragment mass tolerance 0.02, Enzyme: trypsin, Mis-cleavage: −2, Fixed modification: carbamidomethylation of cysteine residues, Dynamic modification: oxidation of methionine and GG modification of lysine. The data were filtered for 1% PSM, peptide level.

The raw data files acquired in DIA mode were processed using DIA-NN, version 1.8 (https://github.com/vdemichev/DiaNN). Human Uniport was used as protein sequence database. Two missed cleavages and a maximum of two variable modifications per peptide were allowed (acetylation of protein N-termini and oxidation of methionine). Carbamidomethylation of cysteines was set as fixed modification and K-GG was added. This data analysis was carried out using library-free analysis mode in DIA-NN by enabling “deep learning-based spectra and RTs prediction”. Match between the runs (MBR) was enabled.

### Visualization of ubiquitin sites on SGK3 and PP2CA structures

We found structures of the 496 residues in human SGK3 (Uniport Q96BR1) to be incomplete, with the only reported X-ray derived structure being the PX domain encompassing residues 10-125 (PDB: 6EDX). Therefore, we generated a model of full length SGK3 using AlphaFold^48^. This model was used for used for displaying ubiquitinated lysine residues with PyMOL Molecular Graphics System v3.0 (Schrödinger, LLC). Rendering of PPP2CA (chain Q) residues 1-309 was performed from the cryo-EM structure of the human Integrator-PP2A complex (PDB: 7CUN).

### Bioinformatic data analysis

For bioinformatic analysis, including volcano plots, results were further processed using Perseus ^49^. The data were loaded in Perseus in .txt format. The replicates were grouped based on their annotation. The data were further transformed in log2 scale. These analysed data were used to generate Volcano plots using GraphPad Prism software (Version 8). We used EnrichR (available at: https://maayanlab.cloud/Enrichr) ^50,51^ to conduct the KEGG pathway analysis, and uncover associations with biological processes, molecular functions, localization data, protein domain architecture information and disease association.

## Data availability

The mass spectrometry proteomics raw data and the corresponding processing reports generated in this study have been deposited to the ProteomeXchange Consortium (http://proteomecentral.proteomexchange.org) via the PRIDE partner repository with the dataset identifier for diGly enrichment optimization (PXD057960), Tau-AdPROM proteome and diGly (PXD059978), PPP2CA degradation and diGly (PXD060403), SGK3-PROTAC1 proteome and diGly (PXD060627).

## Code availability

DIA-NN is freely available for download at https://github.com/vdemichev/diann. Custom python scripts that were used to map di-Gly unique sites between DDA and DIA data sets (Fig. 2B), the code has been deposited to zendo repository https://doi.org/10.5281/zenodo.15489001.

## Lead Contact

Further information and requests for resources and reagents should be directed to and will be fulfilled by the Lead Contact, Gopal Sapkota.

## Author Contributions

GS designed and performed experiments, collected and analysed data and contributed to the writing of the manuscript. SR performed some optimisation experiments. TJM designed the strategies for and generated the CRISPR/Cas9 constructs used in this study. NTW generated some constructs used in this study. GG, LDM, AB and DM analysed the Tau data and provided essential feedback on the manuscript. TKP wrote script for comparing DiGly sites between DDA and DIA runs. MAN assisted with annotating the ubiquitylation residues on PPP2CA and SGK3 protein structures. GPS conceived and supervised the project, analysed data, and contributed to the writing of the manuscript.

## Acknowledgements

GPS is supported by the UKRI Medical Research Council (grant MC_UU_00018/6), VLAIO PRiND (HBC.2019.2939) (awarded jointly to GPS and Johnson & Johnson Innovative Medicine by the Government of Belgium) and the pharmaceutical companies supporting the Division of Signal Transduction Therapy (Boehringer Ingelheim, GSK, Merck-Serono). GS, SR, and GG were supported by VLAIO PRiND (HBC.2019.2939). GS was supported by UKRI MRC grant awarded to GPS for a substantial part of the project (grant MC_UU_00018/6). We thank the Sapkota lab members and Neuroscience Discovery teams at Johnson & Johnson for critical feedback. We thank Dario Alessi and Alessio Ciulli for providing us with SGK3 PROTAC1 for this study. We thank Raja S. Nirujogi for valuable discussions. We thank E. Allen, A. Muir, S. Dalglish, E. Webster and J. Stark for assistance with tissue culture, the staff at the DNA Sequencing service (School of Life Sciences, University of Dundee), and the cloning and antibody teams within the MRC PPU Reagents and Services (University of Dundee, coordinated by J. Hastie). We thank the MRC PPU mass spectrometry service team for their help with the project. Some figures were generated using BioRender.com (license number BP26ZAGDQP).

---

**Figure S1.**
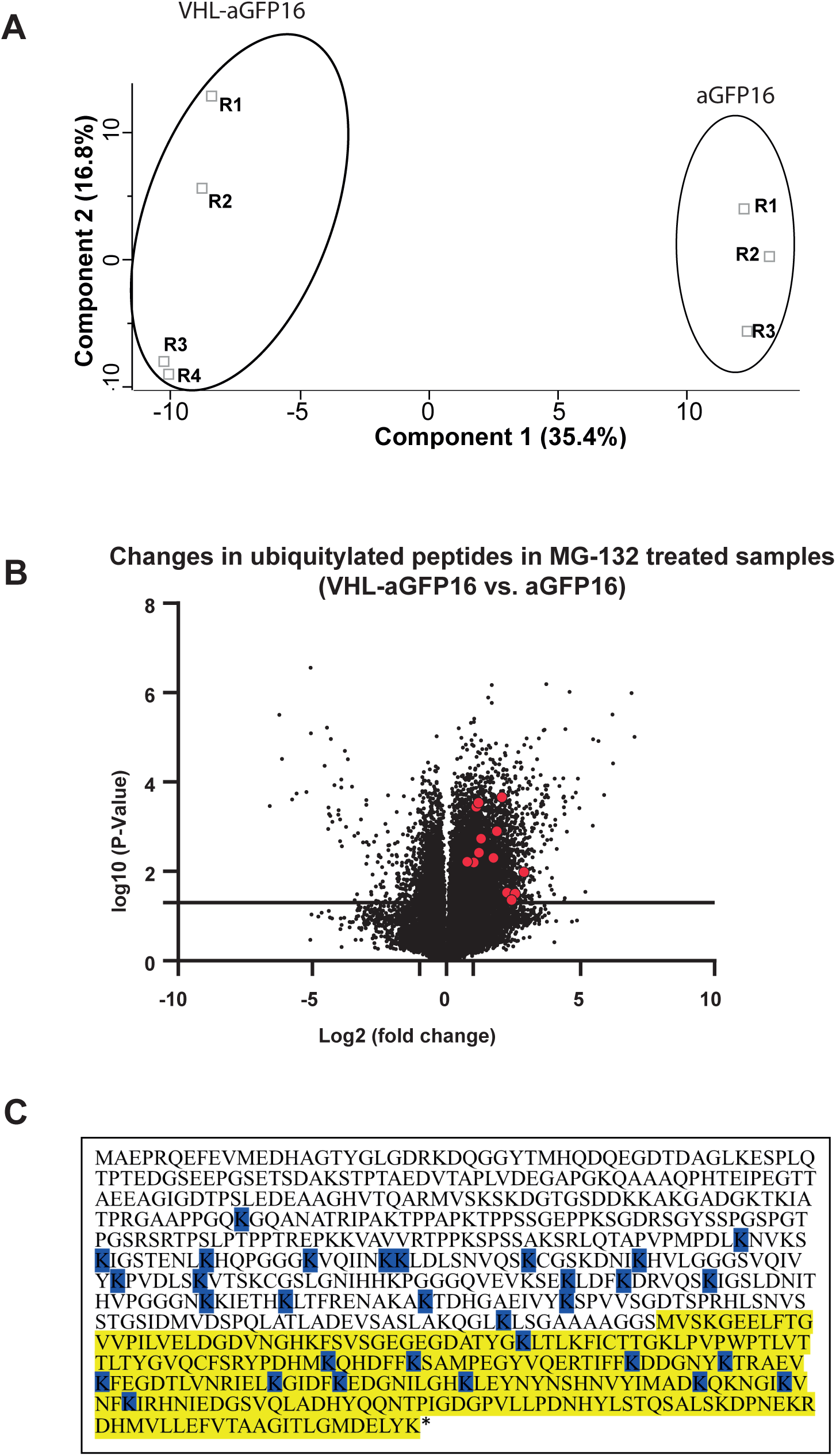
Degradation of Tau-GFP by VHL-aGFP16 AdPROM. **A.** The PCA plot for the proteomics replicate samples for VHL-aGFP16 AdPROM-mediated degradation of Tau-GFP demonstrates that replicates cluster closely together, indicating high reproducibility. Additionally, the two different conditions, VHL-aGFP16 (4 replicates) vs. aGFP16 control (3 replicates), exhibit distinct proteomic profiles, highlighting condition-specific variations. **B.** Volcano plot showing global ubiquitylated peptide changes in extracts of cells expressing VHL-aGFP16 compared to those expressing the aGFP16 control, in the presence of the proteasomal inhibitor MG-132. Indicated in red are some of the significantly ubiquitylated peptides identified on Tau caused by VHL-aGFP16 AdPROM. **C.** All VHL-aGFP16 AdPROM-mediated Tau-GFP ubiquitylation sites are mapped on the amino acid sequence of Tau-GFP. The yellow-highlighted region represents the GFP protein sequence, whereas the remaining sequence corresponds to Tau.

**Figure 2.**
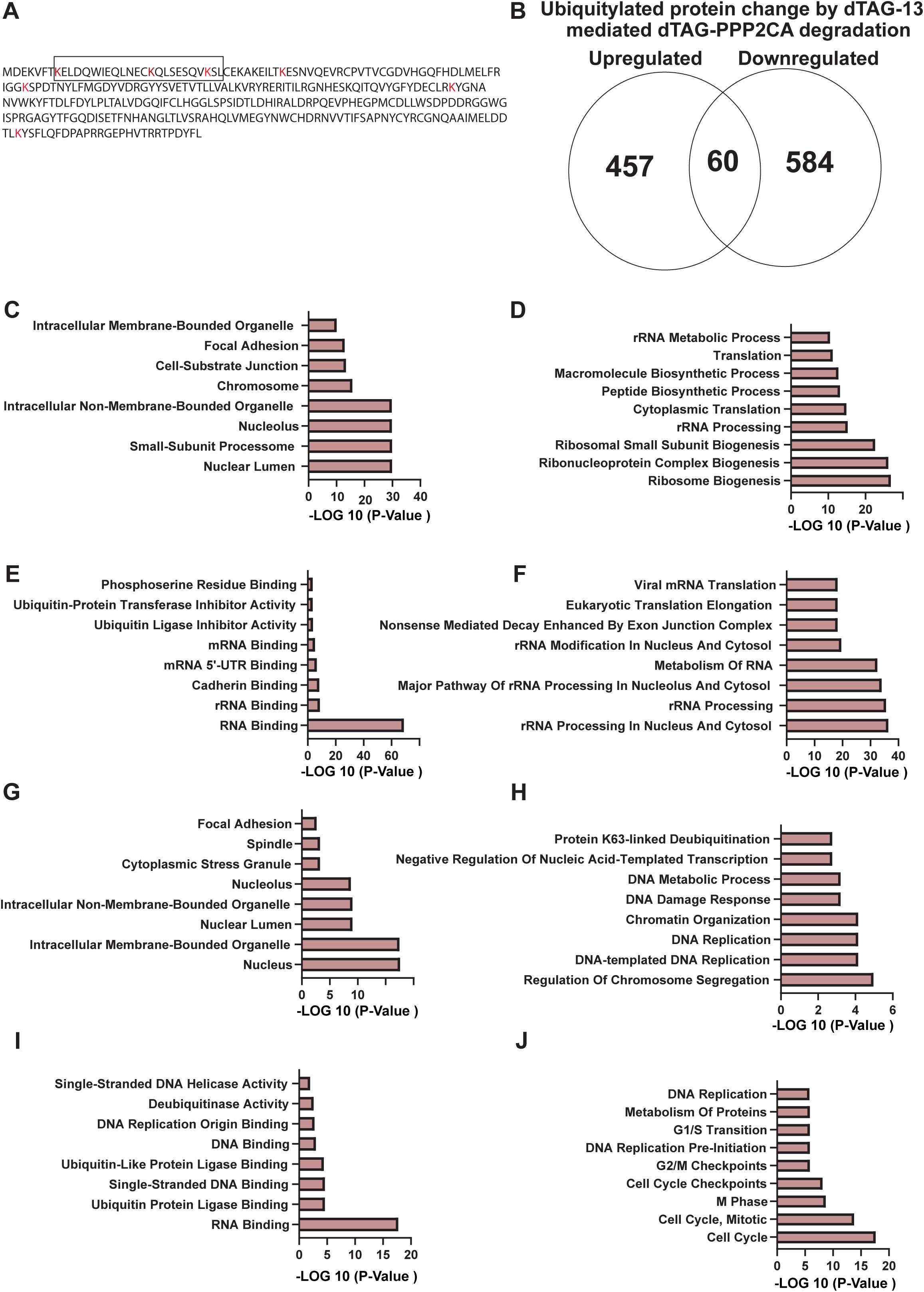
Ubiquitylation of dTAG-PPP2CA upon dTAG-13 treatment. **A.** The amino acid sequence of PPP2CA highlighting all lysine residues that were identified as ubiquitylated upon dTAG-13 treatment. Lysine residues marked red within the boxed region indicate those where higher ubiquitylation was observed at an early dTAG-13 treatment time points. **B.** The Venn diagram shows the number of ubiquitylated proteins that were either up-regulated or down-regulated upon dTAG-13 treatment. The common ubiquitylated proteins reveal that some peptides underwent higher ubiquitylation while other peptides on the same protein showed a reduction in ubiquitylation. **C-G.** GO analysis of the cellular localization (C), molecular function (D), biological function (E) and cellular pathway (F) of proteins that underwent higher ubiquitylation upon degradation of dTAG-PPP2CA by dTAG-13 **H-J.** GO analysis of the cellular localization (C), molecular function (D), biological function (E) and cellular pathway (F) of proteins that underwent lower ubiquitylation upon degradation of dTAG-PPP2CA by dTAG-13

**Figure 3.**
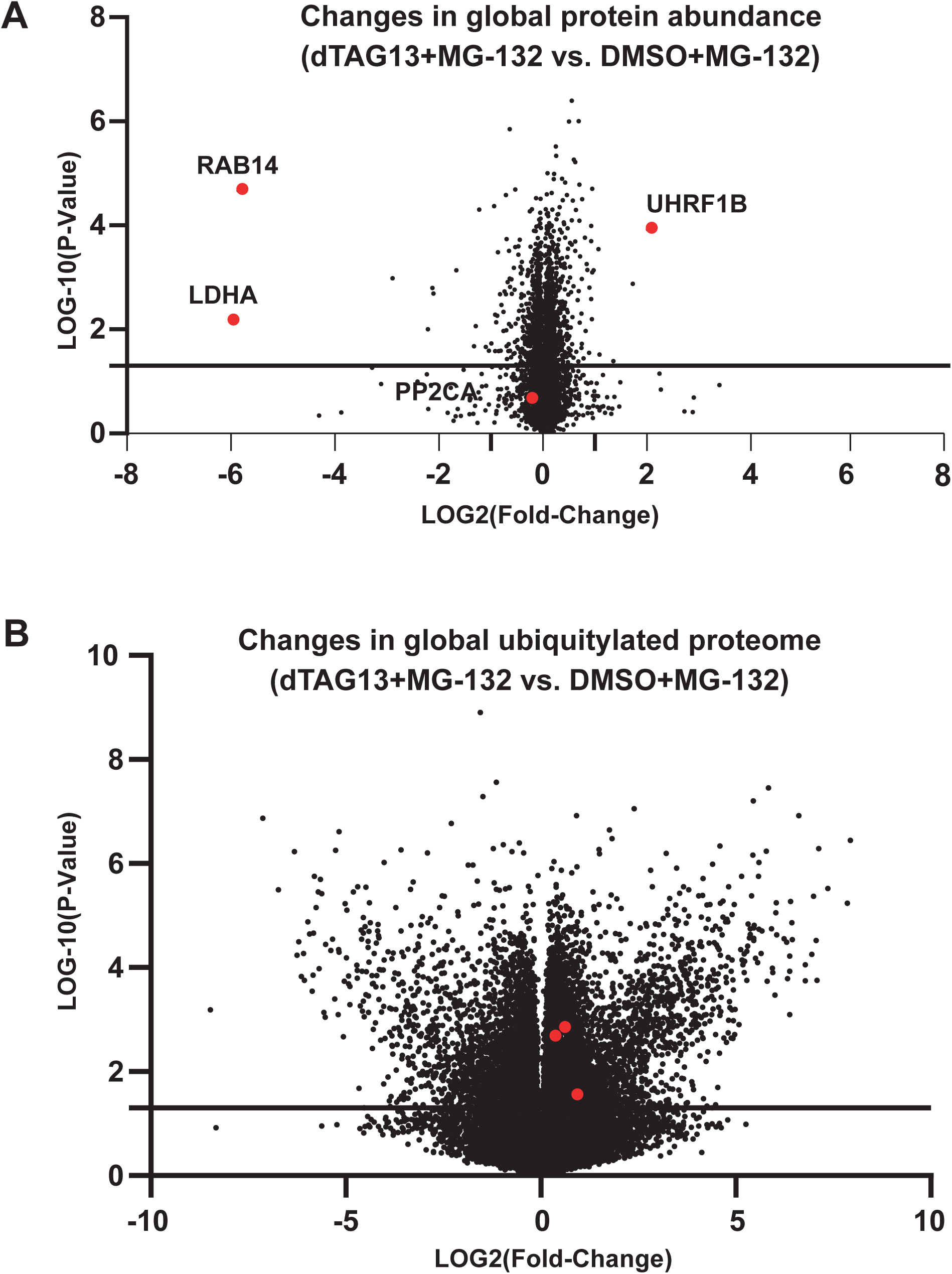
Ubiquitinome changes upon degradation of dTAG-PPP2CA by dTAG-13 in the presence of MG-132. **A.** Volcano plot showing changes in proteins identified from *^dTAG/dTAG^PPP2CA* knock-in HEK293 cells treated with dTAG-13 compared to DMSO in the presence of MG-132. **B.** Volcano plot showing changes in ubiquitylated peptides identified from *^dTAG/dTAG^PPP2CA* knock-in HEK293 cells treated with dTAG-13 compared to DMSO in the presence of MG-132.

**Figure 4.**
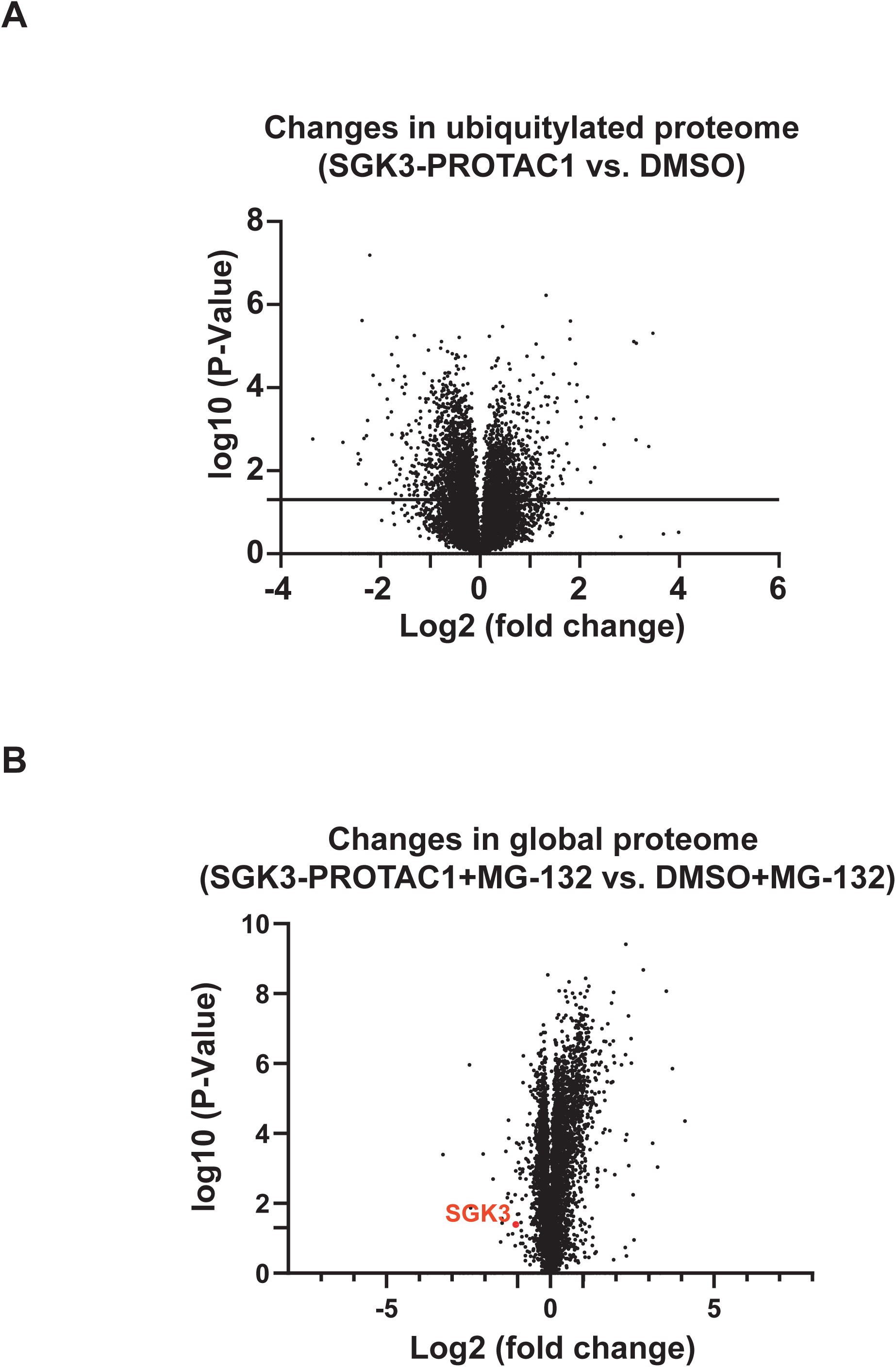
Ubiquitinome and total proteome changes upon treatment of cells with SGK-PROTAC1 in the absence or presence of MG-132. **A.** Volcano plot representing global changes in the abundance of ubiquitylated peptides from HEK293 cells treated with SGK3-PROTAC1 compared to DMSO control in the absence of MG-132. **(B)** Volcano plot showing global changes in the proteome abundance from HEK293 cells treated with SGK3-PROTAC1 compared to DMSO control in the presence of MG-132.

## Supplementary Data

**Data 1.** The data file contains peptide spectrum map (PSMs) corresponding to DiGly peptides identified from two different lysis methods: urea and SDS. Following lysis, proteins were digested and DiGly peptides were enriched using DiGly specific antibody conjugated to agarose beads. Following clean-up steps, these peptides were analysed using data dependant acquisition (DDA) on mass spectrometer and analysed using SEQUEST search engine. The data were obtained for 3 replicates (R1-R3).

**Data 2.** Isolation windows used for data independent acquisition (DIA) method are indicated.

**Data 3.** The data contains the list of DiGly peptides identified from DIA for individual experimental replicate (R1-R3). Cells were lysed in urea lysis buffer, proteins were digested and DiGly peptides were enriched using DiGly specific antibody conjugated to agarose beads. The data were acquired using DIA method and analysed using DIA-NN search engine.

**Data 4.** The data contains the list of DiGly modified proteins identified from DIA for each replicate from Data 3.

**Dats 5.** The data contains the list of DiGly peptides identified from DIA for individual experimental replicate (R1-R3). Cells were lysed in urea lysis buffer, proteins were digested and DiGly peptides were enriched using DiGly specific antibody conjugated to magnetic beads. The data were acquired using DIA method and analysed using DIA-NN search engine.

**Data 6.** As in legend to Data 5, except that the data was analysed using Spectronaut search engine.

**Data 7.** The data contains the list of the identified proteins from total proteomic analysis of extracts from cells expressing VHL-aGFP16 AdPROM compared to those expressing the aGFP16 control.

**Data 8.** The data contains a summary of the DiGly peptides identified through global ubiquitinome analysis of extracts from VHL-aGFP16 AdPROM-expressing cells relative to those expressing the aGFP16 control in Tau-GFP expressing cells upon treatment with MG-132.

**Data 9.** The data contains the list of identified proteins from a total proteomic analysis of extracts from dTAG-PP2CA KI HEK293 cells treated with dTAG-13 compared to DMSO control. Data for three replicates (R1-R3) for each treatment condition is included.

**Data 10.** The data summarizes DiGly peptides identified through global ubiquitinome analysis from extracts of dTAG-PP2CA KI HEK293 cells treated with dTAG-13 compared to DMSO controls in 3 replicates (R1-R3) for each condition.

**Data 11.** The data contains the list of identified proteins from total proteomic analysis from extracts of dTAG-PP2CA KI HEK293 cells treated with dTAG-13+MG-132 compared to DMSO+MG-132 controls in 3 replicates (R1-R3) for each condition.

**Data 12.** The data summarizes DiGly peptides identified through global ubiquitinome analysis from extracts of dTAG-PP2CA KI HEK293 cells treated with dTAG-13+MG-132 compared to DMSO+MG-132 controls in 3 replicates (R1-R3) for each condition.

**Data 13.** The data lists DiGly peptides identified through global ubiquitinome analysis from extracts of dTAG-PP2CA KI HEK293 cells treated with dTAG-13 for 15,30 and 60 minutes compared to DMSO controls in 3 replicates (R1-R3) for each condition.

**Data 14.** The data contains the list of identified proteins from total proteomic analysis from extracts of HEK293 cells treated with SGK3-PROTAC1 compared to DMSO controls in 3 replicates (R1-R3) for each condition in the absence of MG-132.

**Data 15.** The data summarizes DiGly peptides identified through global ubiquitinome analysis from extracts of HEK293 cells treated with SGK3-PROTAC1 compared to DMSO controls in 3 replicates (R1-R3) for each condition in the absence of MG-132.

**Data 16.** The data contains the list of identified proteins from total proteomic analysis from extracts of HEK293 cells treated with SGK3-PROTAC1+MG-132 compared to DMSO+MG-132 controls in 3 replicates (R1-R3) for each condition.

**Data 17.** The data summarizes DiGly peptides identified through global ubiquitinome analysis from extracts of HEK293 cells treated with SGK3-PROTAC1+MG-132 compared to DMSO+MG-132 controls in 3 replicates (R1-R3) for each condition.

